# An ecological-evolutionary perspective on the genomic diversity and habitat preferences of the Acidobacteriota

**DOI:** 10.1101/2024.07.05.601421

**Authors:** Ella McReynolds, Mostafa S. Elshahed, Noha H. Youssef

## Abstract

Members of the phylum Acidobacteriota inhabit a wide range of ecosystems including soils. We analyzed the global patterns of distribution and habitat preferences of various Acidobacteriota lineages across major ecosystems (soil, engineered, host-associated, marine, non-marine saline and alkaline, and terrestrial non-soil ecosystem) in 248,559 publicly available metagenomic datasets. Classes Terriglobia, Vicinamibacteria, Blastocatellia, and Thermoanaerobaculia were highly ubiquitous and showed clear preference to soil over non-soil habitats, class Polarisedimenticolia showed comparable ubiquity and preference between soil and non-soil habitats, while classes Aminicenantia and Holophagae showed preferences to non-soil habitats. However, while specific preferences were observed, most Acidobacteriota lineages were habitat generalists rather than specialists, with genomic and/or metagenomic fragments recovered from soil and non-soil habitats at various levels of taxonomic resolution. Comparative analysis of 1930 genomes strongly indicates that phylogenetic affiliation plays a more important role than the habitat from which the genome was recovered in shaping the genomic characteristics and metabolic capacities of the *Acidobacteriota*. The observed lack of strong habitat specialization and habitat transition driven lineage evolution in the Acidobacteriota suggest ready cross colonization between soil and non-soil habitats. We posit that such capacity is key to the successful establishment of Acidobacteriota as a major component in soil microbiomes post ecosystem disturbance events or during pedogenesis.

## Introduction

The phylum Acidobacteriota [1] (previously, the Acidobacteria) represents one of the most prevalent phyla encountered in soils, as evident in 16S rRNA gene-based surveys, isolation efforts, and metagenomic studies [2–15]. Additionally, members of the phylum Acidobacteriota are also encountered in a wide range of habitats other than soil, e.g. hydrothermal vents [16], anoxic freshwater mud [17], hydrocarbon-contaminated aquifer [18], marine chiton microbiome [19], alkaline hot springs [20], termite nests [21], and marine sediments [22]. Several classification schemes have been proposed for the Acidobacteriota [2, 23, 24]. The Genome Taxonomy Database (GTDB, release 214) [25] classifies the phylum Acidobacteriota into 15 classes, 54 orders, and 102 families, many of which do not have a pure culture representative.

Based on isolation and culture-independent efforts, preference of specific Acidobacteriota lineages to a certain habitat has been observed. For example Acidobacteriota groups 1, 3, 4, and 6 (Barns [2] classification scheme) are the most prevalent members of the Acidobacteriota in diversity surveys [9] with several representative isolates described from soil [6, 7, 26–29], while representatives of groups 8, 10, and 23 have been isolated form non-soil habitats (e.g. hydrothermal vent, chiton microbiome, anoxic sediments, hot springs) [16–18, 30]. However, a detailed global meta-analysis to elucidate patterns of distribution and habitat preference for cultured and yet-uncultured lineages at various levels of phylogenetic resolution is currently lacking. The current availability of hundreds of thousands of metagenomic datasets, as well as a genome-based reference database and taxonomic outlines that encompass cultured as well as uncultured taxa [25] allows for assessing global ecological distribution patterns of target Acidobacteriota lineages at various levels of phylogenetic resolution using metagenomic read-mapping approaches. Such approaches are superior to assessments that rely on documenting the habitat origin of isolates, 16S rRNA sequences, or genomes deposited in databases, where issues regarding differential amenability of lineages to culturing, sequence deposition procedures, preferential amplification of lineages, and genome assembly problems could impact the accuracy of the obtained outcome. As well, read mapping-based assessments would go beyond cursory characterization of habitat preferences by providing quantitative metrics for ubiquity (occurrence of a lineage in a specific ecosystem), preference (comparison of ubiquity levels of a specific lineage across ecosystems), and relative abundance (percentage of reads belonging to a specific lineage within a metagenomic dataset). Further, such assessment could also be used to identify whether a specific lineage is a habitat specialist (i.e. restricted to a single habitat) or a generalist (i.e. encountered in a wide range of habitats) at various taxonomic resolutions.

Progress in isolation, genome sequencing and -most importantly-generation of single cell genomes (SAGs) and metagenome-assembled genomes (MAGs) from environmental samples has greatly increased the number of publicly available Acidobacteriota genomes. Currently, the genome taxonomy database (GTDB) contains 2028 Acidobacteriota genomes from a wide array of habitats. As such, a phylum-wide comparative genomic analysis to elucidate and expand on key genomic features, metabolic capacities, and physiological preferences within various members in the Acidobacteriota is feasible. Such detailed analysis could also be used to disentangle the relative importance of phylogeny (lineage to which a genome belongs) versus habitat (origin from which the isolate, MAG, or SAG was recovered) in shaping genomic features and metabolic capacities in the Acidobacteriota.

Here, we present a detailed analysis of the global distribution patterns of the Acidobacteriota based on fragment recruitment from 248,559 metagenomic studies. We combine this analysis with a detailed comparative genomic analysis based on thousands of genomes available in GTDB to identify key differences across lineages and habitats. Based on our results, we propose the occurrence of ready cross colonization of Acidobacteriota between soil and non-soil habitats through a continuous niche-selection process. We posit that such capacity is key to the rapid and successful establishment of Acidobacteriota as a major component of the soil microbiome during pedogenesis or during recolonization post drastic disturbance events. The implications of such findings on our understanding of how the evolution of soil as a distinct habitat on earth impacted the evolutionary trajectory of the *Acidobacteriota* is further discussed.

## Materials and Methods

### Taxonomic framework for the Acidobacteriota

Several classification schemes have been proposed for the phylum Acidobacteriota. Barns et al [2] classified the phylum into 26 subgroups based on amplicons identified in subsurface sediments. Dedysh and Yelmaz built on this scheme and assigned the 26 subgroups into 15 class-level units [23]. The Genome Taxonomy Database (GTDB) [25] incorporates genomes from isolates as well as single amplified genomes (SAGs) and metagenome-assembled genomes (MAGs) into a global genome-based taxonomy, while using validly described names for isolates. The GTDB (release 214) classifies the phylum Acidobacteriota into 14 classes, 52 orders, 102 families, and 486 genera. Here, given the need for genome-based analysis, and the fact that the GTDB incorporates genomes from both cultured and uncultured lineages, we used the GTDB as our classification framework. Table S1 and [31] provide a comprehensive view of the phylum classification based on the three schemes discussed above [3, 23, 25], as well as the Silva classification database [24].

### Ecological distribution of the Acidobacteriota

Identification of the taxonomy and relative abundances of key taxa in metagenomic datasets is key to deciphering their ecological distribution. We used Sandpiper, an interface that utilizes a recently developed tool (SingleM) for accurate mapping of metagenomic reads to genomes, to determine the occurrence and relative abundance of the phylum Acidobacteriota, its classes, orders, and families [32] in metagenomes. Our search comprised 248,559 metagenomic datasets in 17,617 projects with 1.3 Pbp of sequencing data. For each taxon searched, the output includes the number of datasets where the taxon was identified, along with individual accession numbers and ecological classification of each dataset, as well as relative abundance of the taxon in each dataset. We believe that this approach is superior to other approaches for metagenomic profiling, since shotgun sequencing directly from an environment provides an unbiased view of the available community that is not prone to experimental issues, e.g. primer bias, that would be encountered with culture-independent studies. Also, searching metagenomic reads obtained with shotgun sequencing to identify taxa without assembling the reads into contigs and binning contigs into genomes (MAGs) would provide access to the fraction of the community that would otherwise not be binned into MAGs due to variability in coverage.

We used the output from Sandpiper searches to assess the distribution and prevalence patterns of the phylum Acidobacteriota, as well as its classes, orders, and families recognized in the GTDB taxonomy across six major habitats, as defined by [33]: soil (n=10,250 datasets), engineered (n=16,320 datasets), host-associated (n=165,152 datasets), marine (n=13,880 datasets), non-marine saline and alkaline ecosystems (n=1306 datasets), and terrestrial non soil ecosystems (n=5039 datasets). For each taxon (the phylum, as well as each class, order, family), three criteria were assessed: (1) ubiquity: assessed as the percentage of the total datasets from each habitat classification where a specific lineage was identified by read mapping, (2) Preference of a specific lineage for soil: assessed as the ratio of soil/non-soil datasets where a specific lineage was identified by read mapping, and (3) relative abundance of a specific lineage: assessed as the percentage of reads in a specific dataset that mapped to a target lineage. Due to the overrepresentation of host-associated (mostly human and murine) datasets in that database, and the well-established scarcity of Acidobacteriota in this habitat, all comparative analysis between soil and non-soil habitats, e.g. preference values, were conducted after exclusion of the host-associated datasets.

### Habitat generalization versus specialization estimation

Habitat specialization was assessed by identifying the range of habitats where a specific lineage (class, order, family) was encountered. The analysis was conducted for all taxa (classes, orders, and families) encountered in the GTDB database. However, it is important to note that taxa rarely encountered could erroneously be classified as specialists. Therefore, we also re-analyzed such patterns using anempirical cutoff that excludes taxa encountered in less than 250 studies.

### Comparative genomic analysis

We assessed different genomic features, secretory arsenals, predicted physiological preferences, and lifestyle within Acidobacteriota genomes. We compared these characteristics across various lineages, as well as across habitats in a lineage-agnostic manner (i.e. comparing all genomes recovered from soil to those recovered from non-soil habitats). In addition, we also assessed within lineage habitat effects, i.e. whether genomic features differ between the same lineage genomes recovered from soil versus non-soil habitats. Out of the 2028 Acidobacteriota genomes available in GTDB, we focused our comparative genomic analysis on 1930 genomes belonging to classes with at least 100 genomes and for which genomes were recovered from at least 5 of the 7 habitats (Figure S1). These classes are Aminicenantia, Blastocatellia, Holophagae, Terriglobia, Thermoanaerobaculia, and Vicinamibacteria.

General genomic features (including genome size, GC content, coding density, and number of genes) were retrieved from the GTDB metadata file (available at https://data.gtdb.ecogenomic.org/releases/release214/214.1/). Average gene length was calculated using a perl script. CRISPRs were predicted using CCTyper [34]. Viral contigs were predicted using Virsorter2 [35]. For secretory capacities: CAZymes were predicted using dbCAN3 [36], while biosynthetic gene clusters (BGCs) were predicted using antiSMASH [37]. Proteases were identified via comparisons to the MEROPS database [38].

We used recently developed machine-learning (ML) based bioinformatic approaches to predict physiological preferences from genomic data. To predict optimal growth temperatures, we used the Tome suite [39]. pH preferences were assessed using a newly devised ML approach [40]. To predict oxygen preference from genomic data, we used the recently developed aerobicity ML predictor [41].

To predict life history strategy (Scarcity, Ruderal, or Competitor), we used the recently developed machine-learning approach that measures the tradeoff between genomic investment in resource acquisition and regulatory flexibility to predict the ecological strategy [42]. Genomic investment in regulatory flexibility was calculated as the number of transcription factors relative to total gene number. Transcription factors were predicted using two approaches: (1) Examining the output of BlastKOALA [43] for the presence of transcription factor Kegg Orthologies (KOs) (a list is available at https://www.genome.jp/brite/ko03000), and (2) additional transcription factors were predicted using DeepTFactor [44]. Genomic investment in resource acquisition was calculated as the number of genes encoding secreted enzymes (CAZymes, proteases, and lipases/hydrolases) plus the number of BGCs divided by the total number of membrane transporters. CAZymes and proteases were predicted as detailed above. Lipases/hydrolases were identified through comparison to the ESTHER database [45]. All three groups of enzymes were then subjected to SignalP [46] analysis to identify those with a secretion signal. Finally, membrane transporters were predicted by examining the output of BlastKOALA [43] for the presence of transporters Kegg Orthologies (KOs) (a list is available at https://www.genome.jp/brite/ko02000). Both genomic investment in resource acquisition (the number of genes encoding secreted CAZymes, proteases, and lipases/hydrolases plus the number of BGCs divided by the total number of membrane transporters) and in regulatory flexibility (the number of transcription factors divided by the total number of genes) were then used to predict the life history strategy (one of Scarcity, Ruderal, or Competitor) of all Acidobacteriota genomes via kmeans clustering using the R package flexclust [47] and the k-centroid cluster analysis. The training dataset was comprised of the 27 ^13^C-labeled MAGs from [42].

Metabolic potential encoded in Acidobacteriota genomes was predicted using METABOLIC [48].

### Statistical analysis

Analysis of variance (ANOVA) (run through the aov command in R) was used to test for the effect of phylogeny (Class), habitat (the environmental source from which the genomes were obtained), and the interaction between the two on general genomic features (genome size, GC content, coding density, number of genes, and average gene length), phage infection/immunity features (number of viral contigs, number of CRISPR occurrence in a genome), potential extracellular products arsenal (CAZymes, peptidases, BGCs), predicted physiological optima (temperature, pH, and oxygen-preferences), and predicted life history strategy (ruderal, competitor, scarcity). Factors with F-test p-value <1x10^-5^ were considered significant. The % contribution of phylogeny (Class), and habitat (the environmental source from which the genomes were obtained) was calculated based on the F-test sum of squares. For all significant comparisons, the TukeyHSD command in R was used for multiple comparisons of means.

To test for the effect of phylogeny and habitat on metabolic features predicted in the genomes, we first converted the dichotomous output (Presence/absence) from METABOLIC into 1/0 numerical output (if a pathway was identified as present in a genome, a value of 1 was used, while a value of 0 was used if the pathway was absent). Following, the effect of phylogeny and habitat on the metabolic potential (either the pattern of occurrence of KEGG modules or TIGRfam/Pfam/custom HMM functions as predicted by METABOLIC) was tested using ANOVA as explained above for genomic features.

## Results

### Global patterns of Acidobacteriota distribution across biomes

At the phylum level, Acidobacteriota showed the highest level of ubiquity in soil ecosystems, with fragments mapped to 95.7% of soil-derived datasets (9814 out of 10,250 soil datasets) (Table S2, Figure 1A). In comparison, Acidobacteriota-associated metagenomic fragments were mapped to only 37.6% of non-soil-derived datasets (18,027 out of 47,953 datasets from engineered, freshwater, marine, non-marine saline and alkaline, and terrestrial non-soil habitats) (Table S2, Figure 1A). Similarly, Acidobacteriota exhibited higher mean relative abundance in soil-derived datasets, where it was encountered on average in 15.79% of reads, compared to only 3.71% of reads in non-soil-derived datasets (Table S2, Figure 1C).

**Figure 1.**
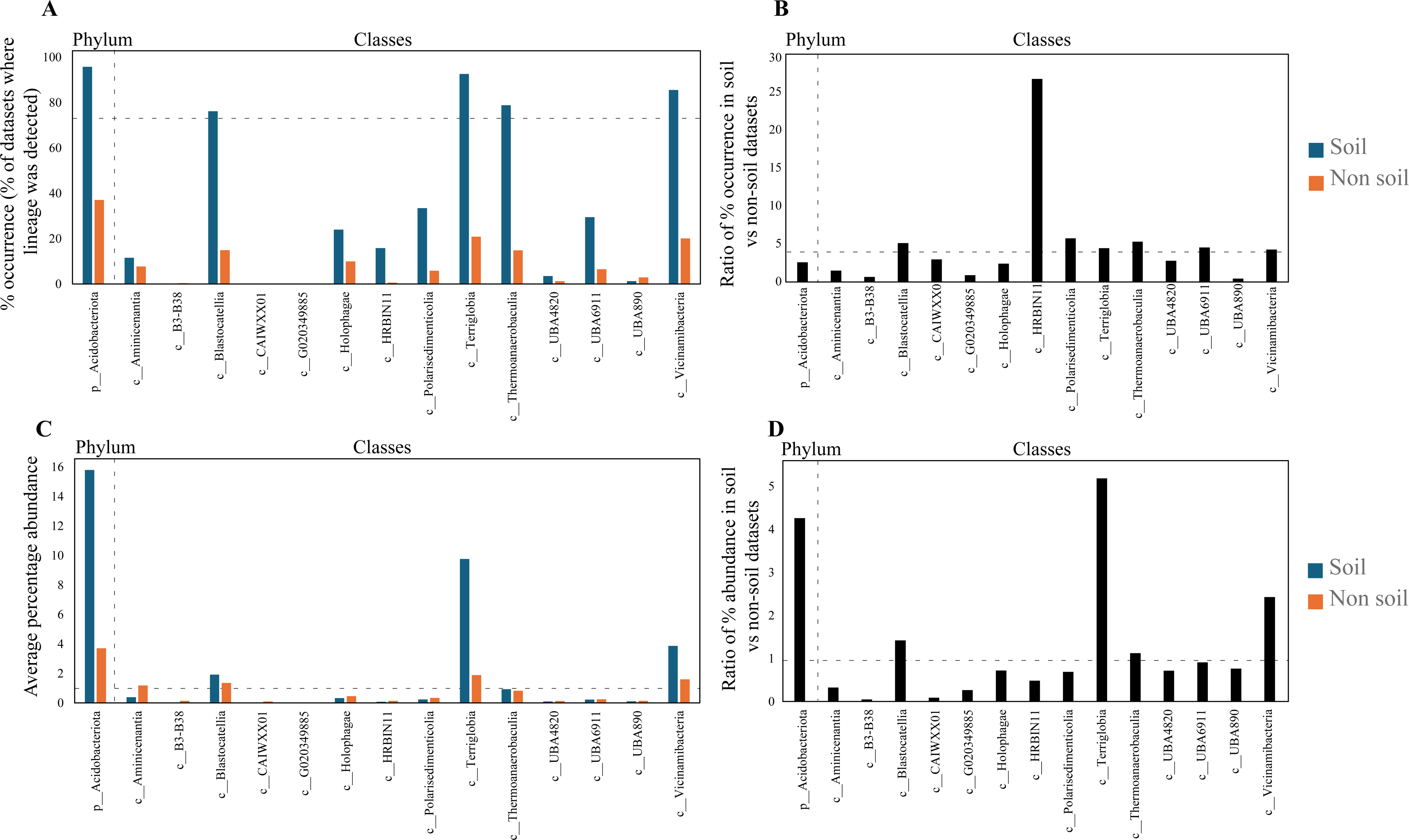
Ecological distribution of the phylum Acidobacteriota and its 14 classes in 248,559 metagenomic datasets available through the web interface Sandpiper [32]. (A) Percentage occurrence of members of the phylum Acidobacteriota and its 14 classes in datasets originating from soil (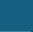), and non-soil (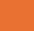) ecosystems. Classes are shown on the X-axis. The dotted line represents 75% occurrence, the cutoff used to define a lineage as ubiquitous in an ecosystem. (B) The ratio of percentage occurrence of the phylum and each of its 14 classes in soil versus non-soil ecosystems. The dotted line represents a ratio of 4, the cutoff used to define a lineage as soil-preferring in an ecosystem. (C) Average percentage abundance of members of the phylum Acidobacteriota and its 14 classes in datasets originating from soil (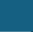), and non-soil (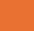) ecosystems. Classes are shown on the X-axis. The dotted line represents 1% occurrence, the cutoff used to define a lineage as abundant in an ecosystem. (D) The ratio of percentage abundance of the phylum and each of its 14 classes in soil versus non-soil ecosystems. The dotted line represents a ratio of 1, the cutoff used to define a lineage as relatively more abundant in an ecosystem.

Four out of the fourteen classes of Acidobacteriota were highly ubiquitous in soil (identified in >75% of soil datasets). These were classes Terriglobia (92.61%), Vicinamibacteria (85.61%), Thermoanaerobaculia (78.84%), and Blastocatellia (76.2%) (Figure 1A, Figure 2, Table S2). In addition, these four classes also showed strong preference to soil (ratio of occurrence in soil to non-soil datasets >4) (Figure 1B, Table S2). This pattern was mostly driven by higher soil ubiquity and preference values for fourteen families belonging to ten orders within these classes (highlighted in Figure 2). Interestingly, while many of these lineages have representative isolates and are currently recognized as prevalent members of the microbial communities in soil, e.g. families *Acidobacteriaceae*, *Koribacteriaceae*, *Bryobacteraceae*, *Pyrinomonadaceae*, and *Vicinamibacteraceae* corresponding to subgroups 1, 3, 4, and 6 (Barns classification system); others represent lineages mostly identified through metagenomic studies, and have, so far, no cultured representatives or recognition in prior amplicon-based [2] and amplicon-and isolates-based [23] taxonomic schemes. For example, within the Terriglobia, in addition to families *Acidobacteriaceae* and *Koribactereaceae* (subgroup 1), and *Bryobacteraceae* (subgroup 3), members of the yet uncultured families 20CM-2-55-15, UBA7541, and SbA1 also showed high levels of ubiquity and preference to soil ecosystems (Figure 2). Within the class Thermoanaerobaculia, named based on earliest isolates from high temperature settings [30], multiple uncultured families showed high levels of ubiquity and preference to soil, e.g. families Gp7-AA8, Gp7-AA6, and UBA5704. Similarly, within the class Vicinamibacteria, in addition to the family *Vicinamibacteraceae* (subgroup 6) [7], the uncultured families Fen-336, 2-12-FULL-66-21, SCN-69-37, and UBA2999 showed high levels of ubiquity and preference to soil. Finally, within the Blastocatellia, while members of the family *Pyrinomonadaceae* (subgroup 4) [6] were ubiquitous in soil, the uncultured family UBA7656 also showed high ubiquity and preference to soil.

**Figure 2.**
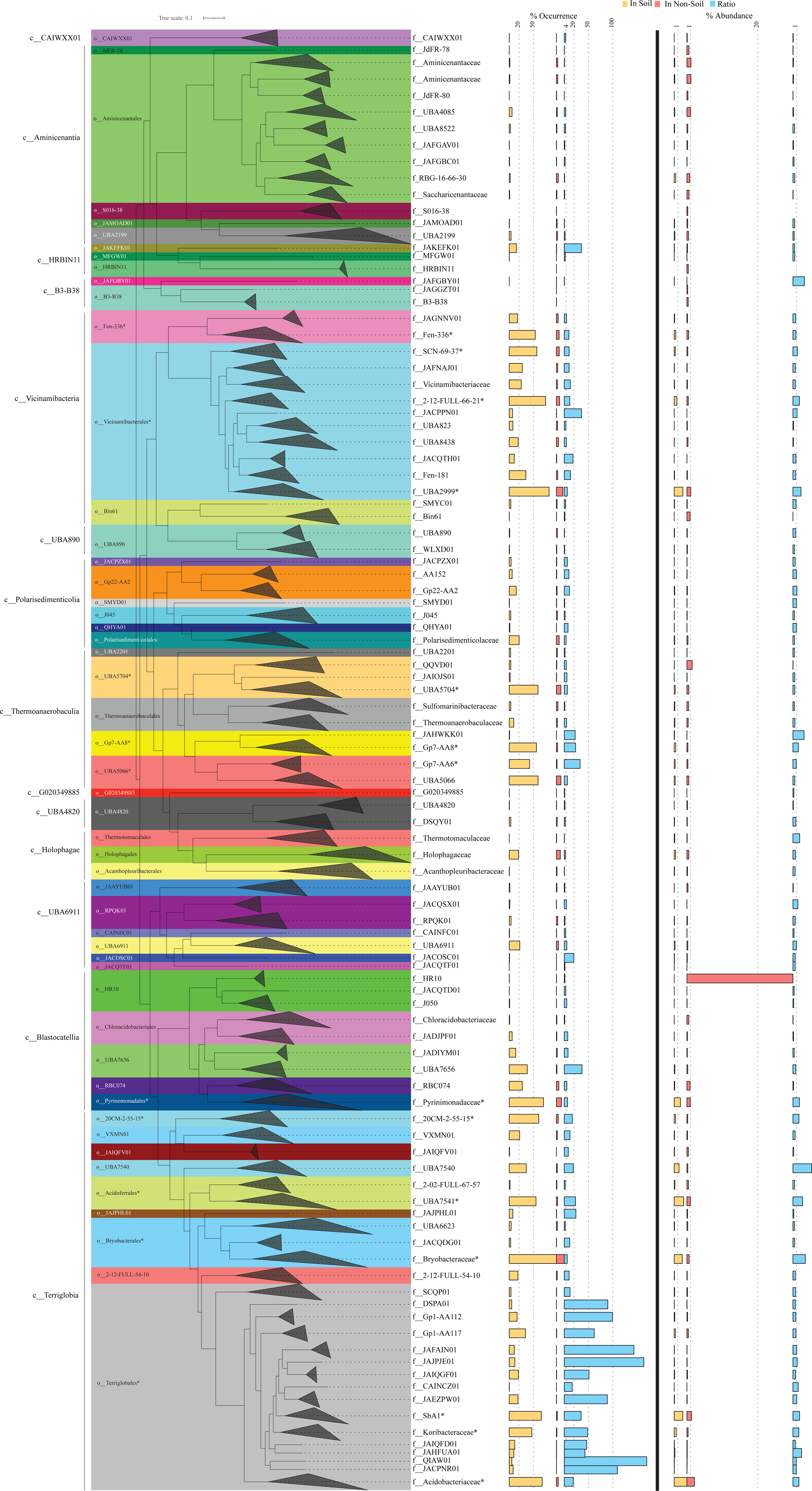
Phylogenomic tree constructed using the GTDB concatenated alignment of 120 single copy marker gene. The tree was constructed in FastTree [71] and is wedged at the family level. Family names are shown on the right. Wedges are color coded by order (shown to the left of the tree). Class names are shown to the left. Families and orders with higher soil ubiquity and preference values are highlighted by an asterisk (*). The values corresponding to the percentage occurrence of each family in soil (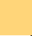), and non-soil (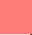) ecosystems, as well as the ratio (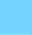) of percentage occurrence in soil versus non-soil ecosystems are shown as horizontal bars to the direct right of the tree. Families with >25% occurrence in soil datasets were considered ubiquitous in soil, while families with a ratio of percentage occurrence in soil versus non-soil ecosystems >4 were considered soil-preferring. For ease of visualization, dashed vertical lines are shown for the 20% and 50% occurrence in soil datasets, and for the soil: non-soil % occurrence ratios 4, 20, 50, and 100. Horizontal bars to the right of the thick vertical line represent the average percentage abundance of each family in soil (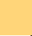), and non-soil (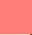) ecosystems, as well as the ratio (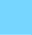) of percentage abundance in soil versus non-soil ecosystems. Families with a ratio of percentage abundance in soil versus non-soil ecosystems >1 were considered more abundant in soil. For ease of visualization, dashed vertical lines are shown for the 1% abundance in soil datasets, 1% and 20% abundance in non-soil datasets, and for the soil: non-soil % abundance ratio of 1.

Acidobacteriota classes that were less ubiquitous in soil included the Polarisedimenticolia (encountered in 33.47% of soil studies), UBA6911 (encountered in 29.48% of soil studies), Holophagae (encountered in 24% of soil studies), HRBIN11 (encountered in 15.78% of soil studies), and Aminicenantia (encountered in 11.5% of soil studies) (Figure 1A, Figure 2, Table S2). Within these five classes, UBA6911, Polarisedimenticolia, and HRBIN11 showed preference to soil (soil: non-soil occurrence ratio >4, Figure 1B, Table S2), while classes Holophagae and Aminicenantia showed lower ratios (<4). Finally, the remaining classes (B3-B38, CAIWXX01, UBA4820, UBA890, and G020349885) were extremely rare (less than 4%) in all habitats examined.

We also assessed relative abundance values (proportion of reads mapped to a specific lineage within a metagenomic dataset) of various classes, orders, and families in the Acidobacteriota, an indirect measure of their niche-colonization capacities and relative contribution to ecosystem functions within a specific habitat (Figure 1C). Classes identified as most ubiquitous in soil also showed the highest relative abundance in soil, with relative abundance values of 9.76%, 3.87%, 1.9%, and 0.92% for Terriglobia, Vicinamibacteria, Blastocatellia, and Thermoanaerobaculia, respectively (Figure 1C). Out of these four classes, Terriglobia had the highest ratio of relative abundance between soil and non-soil datasets (5.2), followed by Vicinamibacteria, Blastocatellia, and Thermoanaerobaculia with soil: non-soil relative abundance ratios of 2.43, 1.42, and 1.12, respectively (Figure 1D). On the other hand, classes with lower ubiquity in soil (Polarisedimenticolia, HRBIN11, Holophagae, Aminicenantia, and UBA6911) showed lower relative abundance in soil compared to non-soil habitats with ratios consistently <1 (Figure 1D, Table S2).

Therefore, based on the above analysis of soil ubiquity, preference, and relative abundance, we infer that members of the four classes Terriglobia, Vicinamibacteria, Blastocatellia, and Thermoanaerobaculia show high level of ubiquity, preference, and relative abundance in soil over other habitats. We refer to these lineages henceforth as “soil-preferring lineages, SPL”. On the other hand, members of the two classes Holophagae and Aminicenantia show lower soil ubiquity, lower preference to soil habitats, and lower relative abundance in soil versus non soil datasets. We refer to these two lineages henceforth as “non-soil preferring lineages, NSPL”. Finally, the three moderately soil-ubiquitous Acidobacteriota classes UBA6911, Polarisedimenticolia, and HRBIN11 showed moderate relative abundance values that were lower in soil, compared to non-soil habitats (values of soil: non-soil relative abundance ratio <1).

### Habitat generalization versus specialization in the Acidobacteriota

We examined patterns of habitat generalization versus specialization in the Acidobacteriota at various levels of taxonomic resolution. We found 78.6%, 61.5%, and 52.9% of classes, orders, and families, respectively, within the Acidobacteriota to be habitat generalists (i.e. with reads belonging to these lineages identified in each of the 7 habitat classifications) (Table S3). However, it is important to note that such pattern could greatly be impacted by sampling depth, with taxa rarely encountered erroneously classified as specialists. Exclusion of rare taxa from our datasets (empirically defined as those encountered in less than 250 studies) results in even higher percentage of generalist taxa (91.7%, 77.5%, and 66.7% of classes, orders, and families, respectively). Our analysis indicate that preferences does not necessarily correspond to specialization, and lineages identified as SPL or NSPL could still be habitat generalists and exhibit wide global multi-habitat distribution patterns. For example, families *Acidobacteriaceae*, *Bryobacteraceae*, *Pyrinomonadaceae*, and *Vicinamibacteraceae* while encountered in 24.76-97.32% of soil datasets, were also present in 2.03-19.75% of engineered, 4-31% of freshwater, 0.61-4.93% of marine, 0.53-10.8% of non-marine saline and alkaline, and 1.47-3.65% of terrestrial non-soil datasets. On the other hand, NSPL families *Aminicenantaceae* and *Holophagaceae,* while enriched in non-soil datasets were also present in 1.23, and 19.26%, respectively, of soil datasets.

To confirm the results obtained from the global metagenomic dataset utilized, we examined the habitat generalization and specialization patterns in the collection of 2028 Acidobacterial genomes available in the GTDB database [25]. Our results (Figure S2, Table S3) confirm habitat generalization of the Acidobacteriota at the class, order, and family levels, with 85.7%, 65.4%, and 54.91% of classes, orders, and families, respectively, within the Acidobacteriota with genome representatives obtained from 2 or more habitat classifications (Table S3). 24.69% of genera had genome representatives obtained from more than one habitat, while 75.3% had genome representatives obtained from only one habitat. Further, when removing lineages defined by less than 5 genomes (404 genera, 48 families, 15 orders, 3 classes), 91.67%, 84.49%, 87.04%, and 62.2% of classes, orders, families, and genera, respectively, were not specialized to one type of habitat (Table S3).

### Comparative genomic features across lineages and habitats

We assessed the general genomic features (genome size, GC content, coding density, average gene length, and number of protein-coding genes (Figure 3)), phage infection/immunity features (number of viral contigs, number of CRISPR occurrence in a genome (Figure 4)), potential extracellular products arsenal (CAZymes, peptidases, BGCs (Figure 5)), predicted physiological optima (temperature, pH, and oxygen-preferences (Figures 6-7)), and predicted life history strategy (ruderal, competitor, scarcity (Figure 8)) in the 1930 genomes belonging to six classes with at least 100 genomes and for which genomes were recovered from at least 5 of the 7 habitats (Aminicenantia, Blastocatellia, Holophagae, Terriglobia, Thermoanaerobaculia, and Vicinamibacteria (Table 1, Figure S1, Table S4)). Comparative analysis was conducted to determine whether significant differences could be identified at a phylogenetic level (i.e. between different lineages within the Acidobacteriota), as well as at the habitat level (i.e. between genomes recovered from soil versus genomes recovered from non-soil habitats, regardless of their phylogenetic affiliation), and the relative contribution of phylogeny versus habitat in shaping Acidobacteriota genomes.

**Figure 3.**
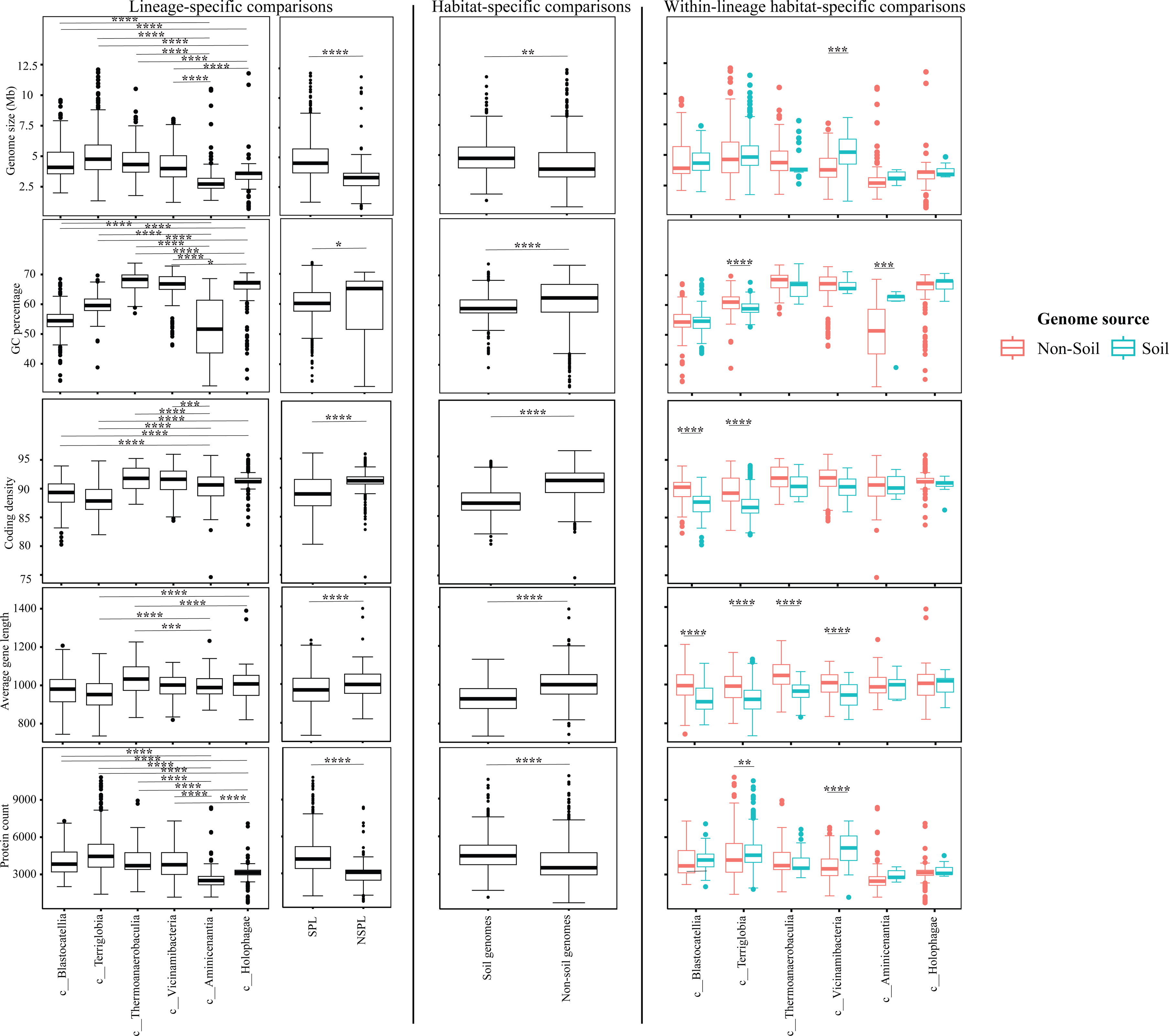
Box plots for the distribution of several general genomic features in the 1930 genomes of the 6 Acidobacteriota classes compared in this study. Features shown are genome size (top row), genome percentage GC (second row from top), genome coding density (third row from top), average gene length (second to last row), and protein count (bottom row). Results for two-tailed ANOVA followed by Tukey for pairwise comparisons are shown on top of the box plots only for significant comparisons. *, 0.01 < p < 0.05; **, p < 0.01; ***, p < 0.001; ****, p < 0.0001; *****, p < 0.00001. The first two columns show results for lineage-specific comparisons at the Acidobacteriota Class level, as well as when combining Acidobacteriota SPL (Terriglobia, Vicinamibacteria, Thermoanaerobaculia, and Blastocatellia) versus NSPL (Holophagae and Aminicenantia). The third column shows results for habitat-specific comparisons (genomes originating from soil versus non-soil environments regardless of phylogeny). Results of within-lineage habitat-specific comparisons are shown in the fourth column, where genomes from soil origin are shown in cyan, while genomes from non-soil origin are shown in red. The exact p-values for all comparisons are shown in Table S4 and S5.

**Figure 4.**
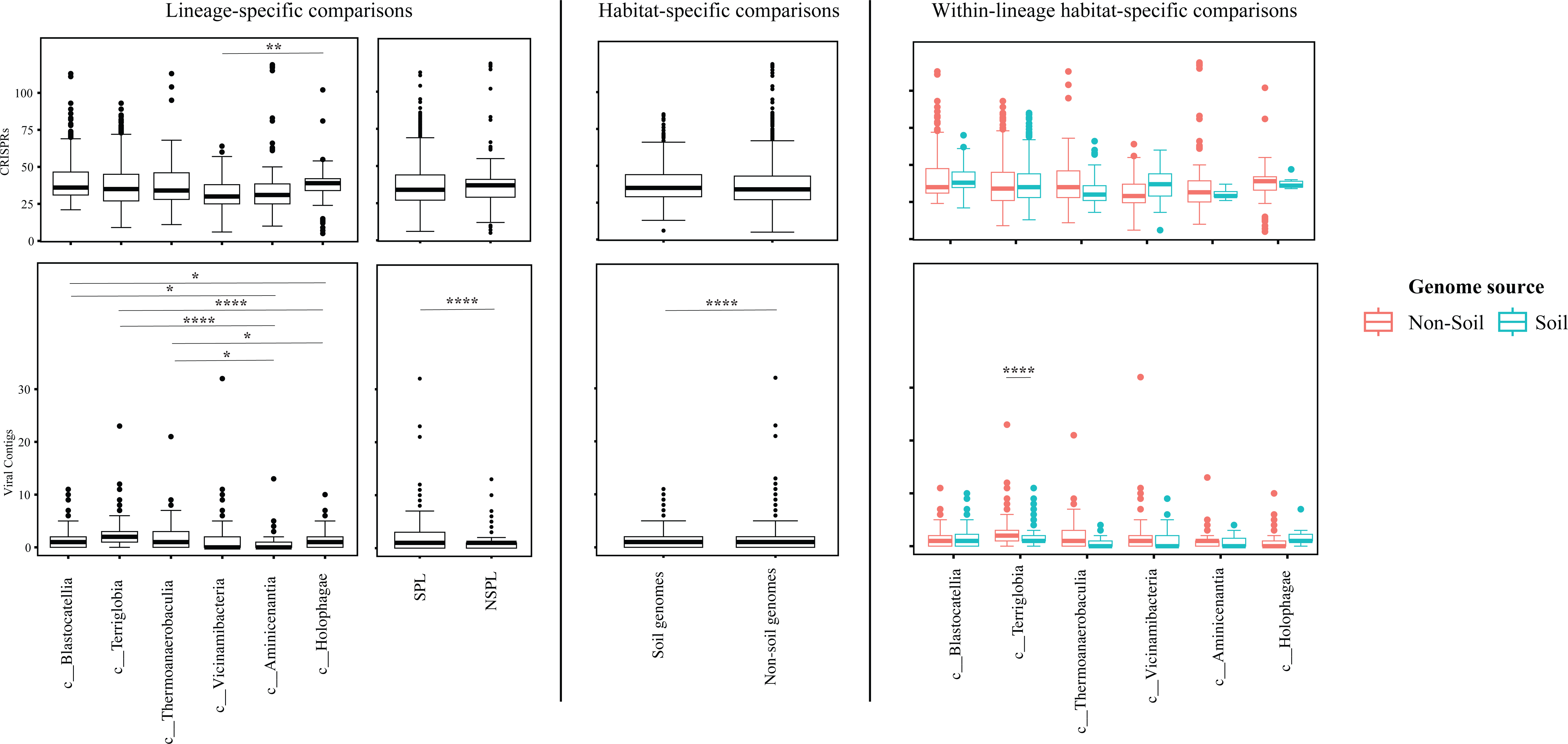
Box plots for the distribution of phage infection/immunity features in the 1930 genomes of the 6 Acidobacteriota classes compared in this study. Features shown are number of CRISPRs (top row), and number of viral contigs (bottom row). Results for two-tailed ANOVA followed by Tukey for pairwise comparisons are shown on top of the box plots only for significant comparisons. *, 0.01 < p < 0.05; **, p < 0.01; ***, p < 0.001; ****, p < 0.0001; *****, p < 0.00001. The first two columns show results for lineage-specific comparisons at the Acidobacteriota Class level, as well as when combining Acidobacteriota SPL (Terriglobia, Vicinamibacteria, Thermoanaerobaculia, and Blastocatellia) versus NSPL (Holophagae and Aminicenantia). The third column shows results for habitat-specific comparisons (genomes originating from soil versus non-soil environments regardless of phylogeny). Results of within-lineage habitat-specific comparisons are shown in the fourth column, where genomes from soil origin are shown in cyan, while genomes from non-soil origin are shown in red. The exact p-values for all comparisons are shown in Table S4 and S5.

**Figure 5.**
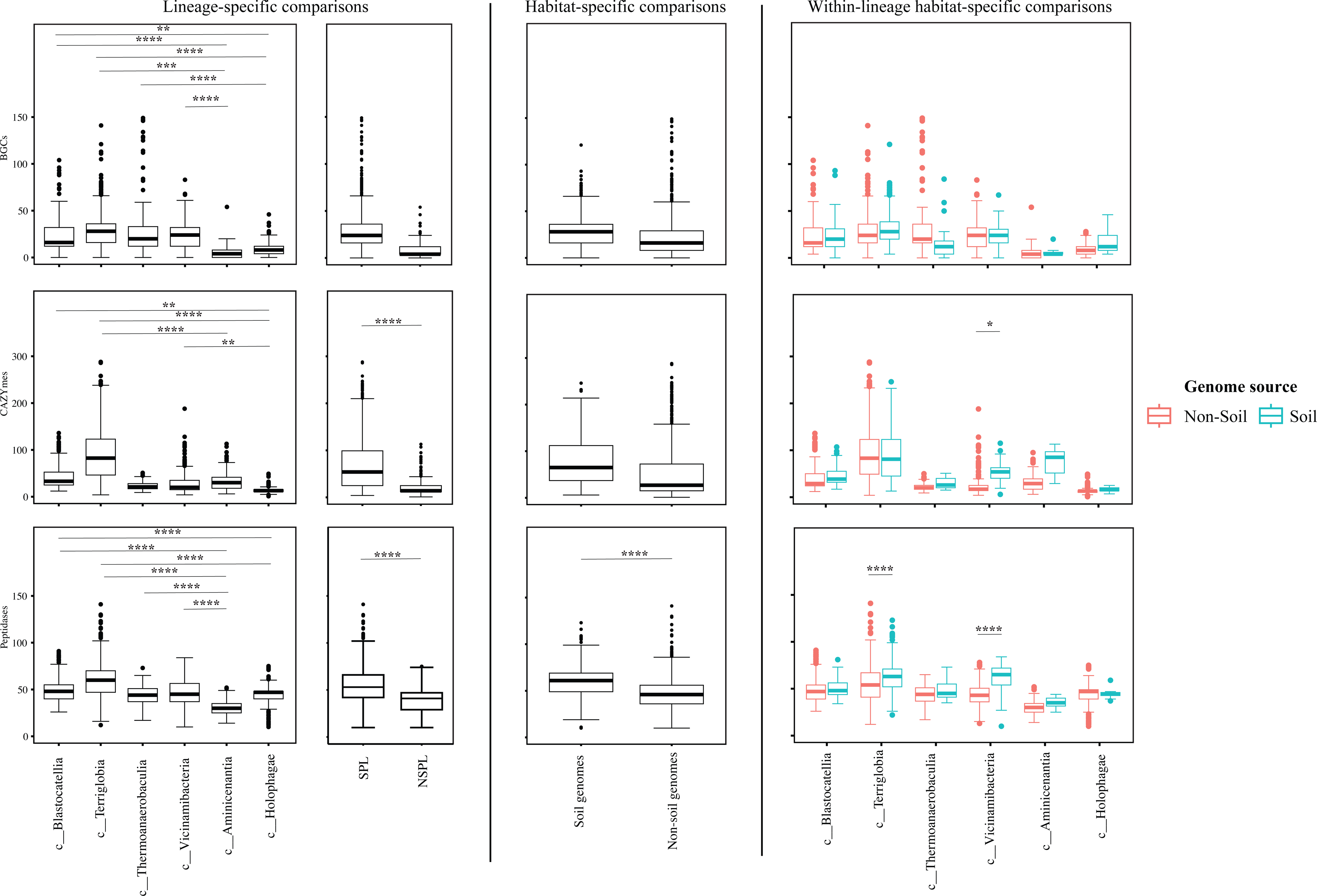
Box plots for the distribution of potential extracellular products arsenal in the 1930 genomes of the 6 Acidobacteriota classes compared in this study. Features shown are number of biosynthetic gene clusters (BGCs, top row), number of CAZymes (middle row), and number of proteases (bottom row). Results for two-tailed ANOVA followed by Tukey for pairwise comparisons are shown on top of the box plots only for significant comparisons. *, 0.01 < p < 0.05; **, p < 0.01; ***, p < 0.001; ****, p < 0.0001; *****, p < 0.00001. The first two columns show results for lineage-specific comparisons at the Acidobacteriota Class level, as well as when combining Acidobacteriota SPL (Terriglobia, Vicinamibacteria, Thermoanaerobaculia, and Blastocatellia) versus NSPL (Holophagae and Aminicenantia). The third column shows results for habitat-specific comparisons (genomes originating from soil versus non-soil environments regardless of phylogeny). Results of within-lineage habitat-specific comparisons are shown in the fourth column, where genomes from soil origin are shown in cyan, while genomes from non-soil origin are shown in red. The exact p-values for all comparisons are shown in Table S4 and S5.

**Figure 6.**
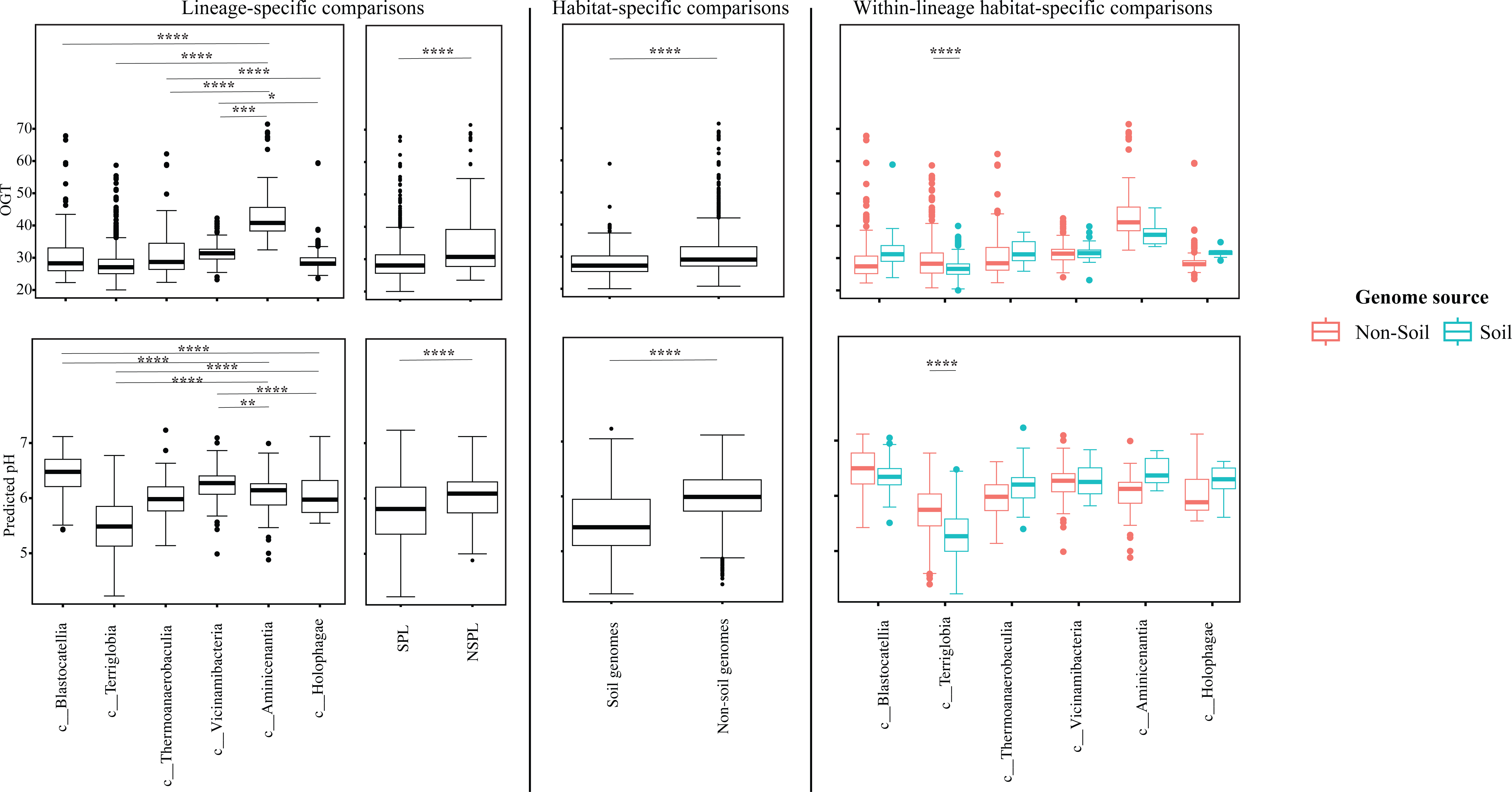
Box plots for the distribution of predicted physiological optima in the 1930 genomes of the 6 Acidobacteriota classes compared in this study. Features shown are predicted optimal growth temperature (OGT, top row), and predicted optimal pH (bottom row). Results for two-tailed ANOVA followed by Tukey for pairwise comparisons are shown on top of the box plots only for significant comparisons. *, 0.01 < p < 0.05; **, p < 0.01; ***, p < 0.001; ****, p < 0.0001; *****, p < 0.00001. The first two columns show results for lineage-specific comparisons at the Acidobacteriota Class level, as well as when combining Acidobacteriota SPL (Terriglobia, Vicinamibacteria, Thermoanaerobaculia, and Blastocatellia) versus NSPL (Holophagae and Aminicenantia). The third column shows results for habitat-specific comparisons (genomes originating from soil versus non-soil environments regardless of phylogeny). Results of within-lineage habitat-specific comparisons are shown in the fourth column, where genomes from soil origin are shown in cyan, while genomes from non-soil origin are shown in red. The exact p-values for all comparisons are shown in Table S4 and S5.

**Figure 7.**
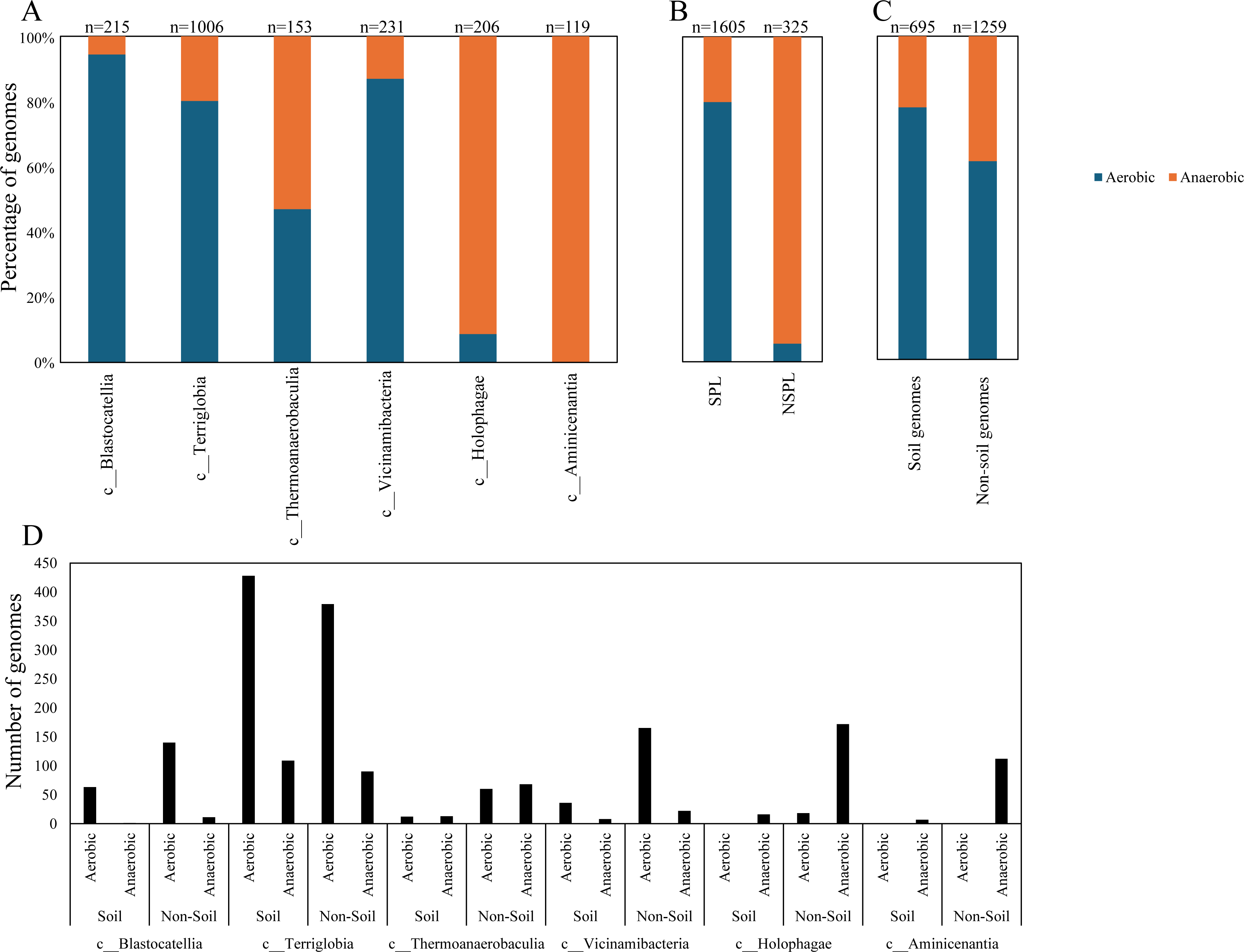
Bar plots for the distribution of the predicted aerobic (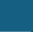) versus anaerobic (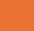) lifestyle in the 1930 genomes of the 6 Acidobacteriota classes compared in this study. Results are shown by class (A), Soil preference (SPL versus NSPL, B), habitat from which the genome originated (C), and within lineage broken down by habitat (D). Number of genomes in each category is shown on top of the bar.

**Figure 8.**
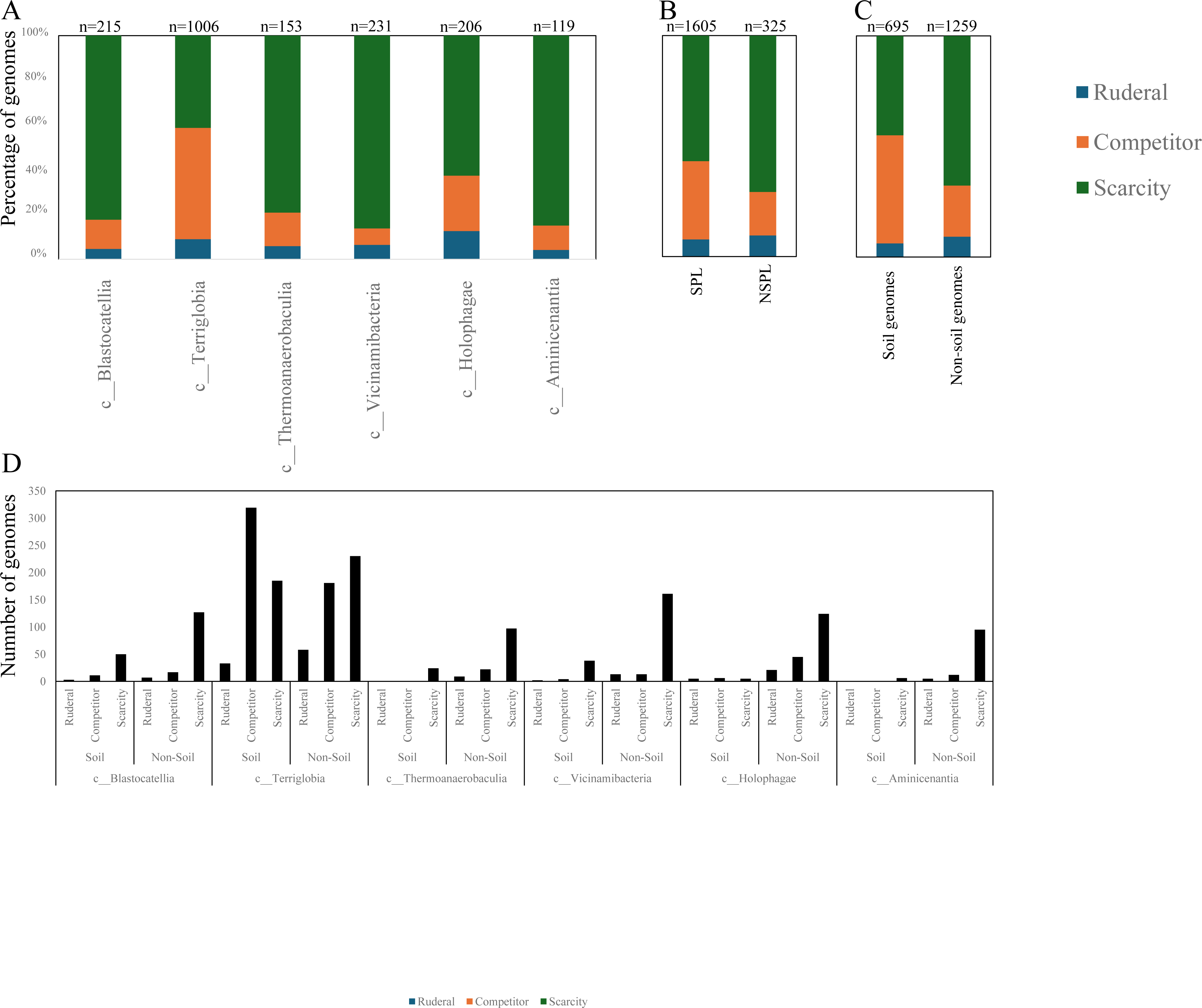
Bar plots for the distribution of the predicted life history strategy (ruderal (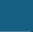), competitor (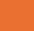), or scarcity (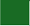) in the 1930 genomes of the 6 Acidobacteriota classes compared in this study. Results are shown by class (A), Soil preference (SPL versus NSPL, B), habitat from which the genome originated (C), and within lineage broken down by habitat (D). Number of genomes in each category is shown on top of the bar.

**Table 1.** Genomes compared in this study. Total number of genomes compared belonging to each of the 6 Acidobacterioat classes is shown. Number of genomes originating from soil, engineered, freshwater, host-associated, marine, non-marine saline and alkaline, and terrestrial non-soil are also shown for each class.

| Class | Total number of genomes | Number of genomes from soil | Number of genomes from non-soil environments |  |  |  |  |  |
| --- | --- | --- | --- | --- | --- | --- | --- | --- |
|  |  |  | Engineered | Freshwater | Host-associated | Marine | Non marine saline and alkaline | Terrestrial non-soil |
| Blastocatellia | 215 | 64 | 103 | 35 | 8 | 1 | 0 | 4 |
| Terriglobia | 1006 | 537 | 179 | 197 | 50 | 16 | 0 | 27 |
| Thermoanaerobaculia | 153 | 25 | 49 | 22 | 37 | 17 | 0 | 3 |
| Vicinamibacteria | 231 | 44 | 35 | 59 | 47 | 46 | 0 | 0 |
| Aminicenantia | 119 | 7 | 27 | 49 | 0 | 35 | 1 | 0 |
| Holophagae | 206 | 16 | 30 | 151 | 4 | 5 | 0 | 0 |

Our analysis demonstrated a significant role played by phylogenetic affiliation in shaping all fourteen examined criteria in Acidobacteriota genomes (F test p-value <1.2 x 10^-10^, percentage lineage contribution 3.6-45.3%, Figures 3-8, Table S5). Interestingly, a trend in some of these criteria was observed where SPL lineages were more similar to each other, compared to NSPL lineages. Specifically, SPL lineages possessed larger genomes (F test p-value = 2x10^-16^, phylogeny percentage contribution=14.96%), higher GC content F test p-value = 2x10^-16^, phylogeny percentage contribution=45.3%), lower gene density F test p-value = 2x10^-16^, phylogeny percentage contribution= 26.31%), shorter average gene length F test p-value = 2x10^-16^, phylogeny percentage contribution= 10.31%), and a higher number of protein-coding genes F test p-value = 2x10^-16^, phylogeny percentage contribution= 18.01%) (Figure 3); as well as a significantly larger number of viral contigs (F test p-value = 4.3x10^-13^, phylogeny percentage contribution= 4.18%) (Figure 4) and a significantly expanded CAZymes (F test p-value = 2x10^-16^, phylogeny percentage contribution= 36.42%), peptidases (F test p-value = 2x10^-16^, phylogeny percentage contribution= 27.83%), and BGCs (F test p-value = 2.6x10^-10^, phylogeny percentage contribution= 3.53%) (Figure 5), and a higher proportion of genes with a predicted aerobic oxygen preference (F test p-value = 2x10^-16^, phylogeny percentage contribution= 41.26%) (Figure 7). For the remaining criteria, while phylogeny played a significant role in shaping the Acidobacteriota genome examined, the SPL lineage: NSPL lineage dichotomy across classes was not observed. For example, while phylogeny had a significant effect on predicted optimal growth temperature (OGT) (F test p-value =2x10^-16^, phylogeny percentage contribution=34.3%), and in general SPL were predicted to have lower optimal pH (Figure 6), this pattern was mainly due to the higher predicted OGT for the Aminicenantia genomes, while Holophagae genomes were predicted to have lower OGT than genomes from SPL lineages (Figure 6). Similarly, while phylogeny had a significant effect on predicted optimal growth pH (F test p-value =2x10^-16^, phylogeny percentage contribution=43.2%), this pattern was mainly due to the lower predicted pH for Terriglobia genomes, while Blastocatellia and Vicinamibacteria genomes were predicted to have higher pH than genomes from NSPL lineages. Finally, for life history strategies (ruderal, competitor, scarcity), phylogeny played a significant role in shaping the predicted life history strategy (F test p-value =2x10^-16^, phylogeny percentage contribution=9.13%), and in general SPL were predicted to have more competitor strategy. However, this pattern was mainly due to the higher number of predicted competitors for Terriglobia genomes, while other SPL classes showed no significant difference in terms of the distribution of predicted life history strategies from the two NSPL classes (Figure 8).

On the other hand, habitat-specific, but lineage-agnostic, comparisons (i.e. comparing genomes originating from soil to those originating from non-soil sources, regardless of their phylogenetic affiliation) identified habitat-specific strong significant difference in 7/14 criteria (F test p-value <10^-5^), with genomes from soil predicted to produce more peptidases, and to have lower coding density, higher number of protein-coding genes, shorter average gene length, lower predicted OGT and lower predicted pH, and predicted to be more competitor. Weak but significant differences were observed for GC percentage (lower in soil genomes), genome size (larger in soil genomes), and number of viral contigs (lower in soil genomes), while the number of predicted BGCs, and CAZymes did not significantly differ by habitat, nor did the total predicted number of CRISPRs, or the preference for O_2_ (Table S5).

While the habitat from which the genome was obtained played a role in shaping genomes in 7/14 criteria; major contribution in these criteria was from the lineage (values of lineage contribution ranging from 3.6-45.3%, as opposed to 0.3-12.3% for the habitat) (Table S5). To further examine differences on a finer scale, we compared within lineages genomes originating from soil versus non-soil habitats. The SPL classes Terriglobia, Vicinamibacteria, Blastocatellia, and Thermoanaerobaculia showed more within-lineage significant differences between genomes originating from soil versus non-soil habitats with 9, 4, 2, and 1, respectively, genomic features identified as within-lineage habitat-specific. These included lower coding density, shorter genes, lower GC percentage, higher number of protein-coding genes, more predicted peptidases, less predicted viral contigs, lower predicted OGT, lower predicted pH, and more predicted competitor lifestyle in Terriglobia soil genomes compared to non-soil genomes, larger genomes, shorter genes, more predicted peptidases and CAZymes in Vicinamibacteria soil genomes compared to non-soil genomes, shorter genes and lower coding density in in Blastocatellia soil genomes compared to non-soil genomes, and shorter genes in Thermoanaerobaculia soil genomes compared to non-soil genomes. On the other hand, NSPL classes showed less within-lineage significant differences between genomes originating from soil versus non-soil habitats with only 1 genomic feature (GC percentage) identified as within-lineage habitat-specific for class Aminicenantia (with soil Aminicenantia genomes having higher GC percentage than non-soil genomes) (Table S5).

### Comparative genomic analysis identifies lineage-specific rather than habitat-specific metabolic differences

We conducted detailed comparative genomic analysis for the metabolic capacities predicted for the 1930 genomes in the SPL classes Blastocatellia, Terriglobia, Thermoanaerobaculia, and Vicinamibacteria, as well as the two NSPL classes Aminicenantia, and Holophagae. We aimed to determine whether significant differences could be identified at a phylogenetic level (i.e. between different lineages within the Acidobacteriota), as well as at the habitat level (i.e. between genomes recovered from soil versus genomes recovered from non-soil habitats, regardless of their phylogenetic affiliation), and the relative contribution of phylogeny versus habitat in shaping Acidobacteriota metabolic capacities.

We identify 73 features where SPL classes differ significantly from NSPL classes (Table S6). These features were mainly distributed among catabolic and anabolic pathways. On the catabolic front, genomes from SPL classes encoded significantly higher O_2_ respiration genes (both low affinity and high affinity cytochrome oxidase), as well as dissimilatory sulfate reduction genes (Table S6). On the other hand, genomes from NSPL classes encoded significantly higher nitrate reduction to ammonium and dissimilatory nitrate reduction genes. Genomes from SPL classes encoded significantly higher genes for ethanol fermentation from acetyl-CoA, while those from NSPL genomes encoded significantly higher phosphate acetyltransferase-acetate kinase pathway for acetate fermentation with concomitant substrate level phosphorylation, and significantly more hydrogenases belonging to the fermentative hydrogenogenic classes [FeFe] hydrogenase group b, [NiFe] hydrogenase group 3b-3d, and [NiFe] hydrogenase group 4a-g, as well as the hydrogenotrophic hydrogenase classes in classes [NiFe] group 1, and [NiFe] group 3c. However, the lack of clear autotrophic potential in these NSPL genomes cast some doubt on these results. It has previously been suggested that atmospheric H_2_ can be scavenged by the thermophilic *Pyrinomonas methylaliphatogenes* K22 in absence of organic carbon as a means of operating the respiratory chain when organic electron donors are scarce [49]. Recently, more Acidobacteriota genomes were shown to encode such hydrogenases [50]. A similar role for these hydrogenotrophic hydrogenases can be predicted for these NSPL classes.

Further, SPL classes were enriched in some central metabolic pathways that are usually associated with the higher O_2_ tension conditions in soil (Table S6). These included the oxidative branch of the PPP, the oxidative TCA cycle, and the semi-phosphorylative Entner-Doudoroff pathway (which joins the oxidative branch of the PPP to glycolysis). SPL genomes are also enriched in the degradation of the sugars/sugar acids galactose, glucuronate, and galacturonate, amino acids Pro, Trp, Met, and hydroxyproline (an amino acid rich in plants cell wall proteins and glycoproteins, and so would be abundant in soil constituting a rich source of C and N for soil bacteria [51]), and in nucleotide degradation (uracil to β-alanine, thymine to 3-amino-isobutanoate, and xanthine to urea) (Table S6).

On the anabolic side, SPL classes exhibited a higher biosynthetic capacity, where biosynthesis of seven amino acids (Lys, Arg, Thr, Val, Ile, His, and Trp) and nine cofactors (Menaquinone, heme, pyridoxal-P, NAD, pantothenate, THF, Lipoic acid, riboflavin, and molybdenum cofactor) were significantly enriched in the four SPL classes (F-test p-value <1e-5). On the other hand, the biosynthesis of only one amino acid (Pro), and two cofactors (thiamine, and biotin) were enriched in the two NSPL classes. Interestingly, some of the cofactors with enriched biosynthesis in the SPL classes are functional during the operation of the electron transport chain for O_2_ respiration (e.g. menaquinone and heme), a trait enriched in SPL lineages (Figure 7). SPL classes were also more enriched in trehalose biosynthesis. The disaccharide trehalose acts as an osmolyte whose biosynthesis was shown to be upregulated in dry soil mainly contributing to tolerance to water deficits [52–55]. On the other hand, NSPL classes were enriched in polyamine biosynthesis. Polyamines are aliphatic polycations with diverse function related to cell growth, biofilm formation, as well as secondary metabolite production [56]. Recently, [57] demonstrated a link between increasing intracellular concentration of polyamines and improved oxidative stress tolerance in *Pseudomonas*. A similar role for polyamine can be speculated in the NSPL classes.

On the other hand, habitat-specific comparisons identified only 45 features where genomes from soil differ significantly from genomes from non-soil sources (Table S6). The majority of these features (42 out of 45) were identified as significantly affected by phylogeny, with only three features (fatty acids beta oxidation pathway genes, N_2_ fixation genes, and high affinity O_2_ respiration using cytochrome oxidase cbb3) identified as habitat-specific (but not lineage-specific). Statistical analysis of features where both phylogeny and habitat were significant (n=42) showed lineage to be more important (lineage percentage contribution: 2-50.8%), with habitat contributing < 7.2%. Of the 42 features where both phylogeny and habitat were significant, 39 followed the same trend (where the feature was higher/lower in SPL as well as genomes sourced from soil regardless of their lineage), with only 3 features with a divergent trend. These later features included phosphatidyl choline biosynthesis, tetrahydrofolate biosynthesis, and nitrite oxidation (significantly higher in SPL genomes and in genomes from non-soil sources).

## Discussion

Here, we examined Acidobacteriota global distribution patterns using a metagenomic read mapping approach, and identified salient differences in genomic characteristics and metabolic capacities across lineages and habitats in the Acidobacteriota. Four classes (Blastocatellia, Terriglobia, Thermoanaerobaculia, and Vicinamibacteria) were shown to have clear preference to soil over non-soil habitats, based on their high soil ubiquity (encountered in >75% of soil datasets), soil preference (ratio of occurrence in datasets from soil versus non soil datasets >4), and higher relative abundance in soil datasets compared to non-soil datasets (Figure 2, Table S2). These four classes encompass many Acidobacteriota taxa previously isolated from soil or shown to occur in high abundance in soil in amplicon-based surveys. The class Terriglobia encompasses subgroups 1 (Families *Acidobacteriaceae* and *Koribacteraceaea*), and 3 (family *Bryobacteraceae*). Class Vicinamibacteria encompasses subgroup 6 (family *Vicinamibacteraceae*). Class Thermoanaerobaculia encompasses subgroups 10, and 23 (family *Thermoanaerobaculaceae*). Class Blastocatellia encompasses subgroup 4 (Families *Pyrinomonadaceae*, *Blastocatellaceae*, and *Chloracidobacteriaceae*). More importantly, we identify multiple additional poorly characterized yet uncultured lineages that are ubiquitous and show high preference and high relative abundance in soil datasets (Figure 2, Table S2). These include the uncultured families UBA7541 (subgroup 2), and SbA1 in Terriglobia, and UBA2999 in Vicinamibacteria. Most of these families are defined based on MAGs recovered from metagenomic studies and are not currently recognized in amplicon-and isolation-based classification. The recognition of the prevalence of such lineages should spur efforts towards more detailed –omics-based characterization and assessment of their importance and potential role in elemental cycling and contribution to ecosystem functioning in soil, as well as stimulate endeavors towards obtaining them in pure cultures.

Organisms encountered in multiple niches are referred to as generalists, while those restricted to a single habitat are specialists. Studies usually utilize occurrence patterns as an empirical measurement of a generalist versus specialist pattern either within a target habitat [58], through a habitat transition gradient, or on a global scale [59]. Assignment of microbial taxa as generalists or specialists could be affected by the level of phylogenetic resolution employed, number of datasets examined, sequencing depth of datasets, and detection threshold employed. Prior research provided some insights on such patterns in the Acidobacteriota. For example, a study examining a large collection of datasets from farmland soils suggested that Acidobacteriota harbored a larger fraction of generalists than specialists at the species OTUs level [58]. On the other hand, studies assessing Acidobacteriota generalist-specialist patterns on a global scale are quite sparse. A recent analysis, defining specialists and generalists using the level of community similarity between datasets where a specific lineage is encountered (with high community similarity indicating a specialist pattern and low community similarity indicating a generalist pattern) as a substitute for occurrence patterns, concluded that Acidobacteriota encompassed a high proportion of specialized genera [59].

Our analysis of Acidobacteriota distribution patterns strongly suggests that a habitat-generalist rather than a habitat-specialist pattern is more common in the Acidobacteriota (Table S3). Using two datasets (publicly available metagenomes via Sandpiper, and publicly available genomes in GTDB), criteria (with and without exclusion of rare taxa, with and without exclusion of taxa with less than 5 genomes), and taxonomic thresholds (class, order, family), we estimate that 73.3-91.7% of classes, 61.5-86.49% of orders, and 52.9-87.04% of families are habitat generalists, with documented ability to inhabit multiple environments.

We attribute the prevalence of habitat-generalist over habitat-specialist pattern in the Acidobacteriota to two possible reasons: the nature of its preferred habitat(s), and the genomic features and metabolic capacities of its members. Broadly, habitats that are extreme, restricted, and drastically different from their surroundings favor specialists, e.g. chemolithotrophic hyperthermophiles in hydrothermal vents [60], strict halophilic Archaea in hypersaline ecosystems [61], anaerobic gut fungi in the herbivorous alimentary tracts [62]. On the other hand, generalists thrive across more temperate, less restricted, and more complex habitats. Broadly, the habitats where Acidobacteriota appears to thrive, e.g. soil, and freshwater, are temperate, with fluctuating temperature, pH, salinity, as well as a complex variable influx of substrates. Such conditions allow for generalists, organisms usually exhibiting a wider arsenal of substrate utilization capacities and response to environmental fluctuations, to thrive as integral components of a complex habitat.

Regarding metabolic abilities, habitat-specialists’ genomes are usually streamlined and either mediate a specific function or encode exceptional capacity to survive and adapt to a unique ecosystem [63]. Habitat-generalists’ genomes, on the other hand, are more metabolically versatile to enable adaptation, e.g. *Pseuodomonas* [64], *Burkholderia* [65] thriving under so many substrates, *Shewanella* thriving under many electron acceptors [66]. Within the Acidobacteriota, genomic analysis identified moderate to large genomes and suggested a general prevalence of heterotrophy, and lack of auxotrophies. Our machine-learning-based analysis suggested prevalence of moderate predicted optimal growth temperature and pH, as well as variable predicted oxygen preferences, all of which would explain the prevalence of habitat-generalist patterns in the phylum. Prior genomic analysis of members of the Acidobacteriota has been conducted on pure culture isolates [14, 50, 67, 68] or assembled MAGs [4, 22, 31, 69, 70]. These efforts have yielded valuable insights into the salient genomic features of various members of the phylum. Our analysis aimed for a broader and more comprehensive view of Acidobacteriota genomes via a global comparative analysis approach. In addition to expanding on known capacities within the phylum, we aimed to identify variations in genomic features, physiological optima, and metabolic capacities within members of the Acidobacteriota, and to determine whether the observed differences are habitat-specific (e.g. present in all genomes of soil Acidobacteriota regardless of the lineage they belong to), lineage-specific (e.g. present in all genomes of a lineage regardless of the environmental source), or a combination of both. Our results suggest that lineage is more important than habitat in describing the observed differences, with very few exceptions. Within general genomics features and physiological optima prediction, lineage significantly affected all 14 criteria compared (F test P-value <2.6 x10^-10^), while habitat only significantly affected 7 criteria, and the interaction between the two only significantly affected 4 criteria (GC percentage, number of predicted peptidases, predicted optimal growth temperature, and predicted optimal growth pH) (Table S5, Figures 3, 5, 6). In all these comparisons, lineage explained 3.5-45.3% of the variability, while habitat only explained 0.91-12.34%. Detailed analysis of metabolic identified 73 functions that were significantly different between SPL and NSPL classes, while habitat only significantly affected 45 functions (42 of which were also significantly affected by lineage). In all these comparisons, lineage explained 2.02-50.8% of the variability, while habitat % contribution never exceeded 7.2%. Indeed, many of the metabolic features previously shown to underpin success of Acidobacteriota in soil, were found to be lineage-specific (significantly encountered more in SPL genomes), rather than habitat-specific (present in all soil Acidobacteriota regardless of their phylogeny). For examples genes encoding CAZymes and BGCs were significantly higher in SPL genomes, but these numbers did not significantly vary in genomes derived from soil versus genomes derived from non-soil environments. However, while clearly lineage plays a more important role, some habitat-specific features are of note. Soil as a habitat appears to select for genomes with lower coding density, higher number of protein-coding genes, shorter average gene length, larger number of peptidases, and for organisms with lower predicted OGT and lower predicted pH, and with a predicted competitor lifestyle.

Finally, the observed prevalence of a generalist patten and higher importance of lineage compared to habitat in shaping Acidobacteriota genomes strongly suggest a ready cross-colonization of Acidobacteriota across major habitats. Under this scenario, while differential preference of certain Acidobacteriota lineages to specific habitat exists, the presence of a ready reservoir, albeit in minor quantities, in other environments would allow ready cross-colonization when needed, e.g. as a mechanism for repopulation post-disturbances (e.g. fire, extensive pollution), or during the process of pedogenesis (soil formation). Nevertheless, our results also advocate for a role played by the environment, specifically soil studied here, in shaping Acidobacteriota genomes post-acquisition. Specifically, larger number of encoded peptidases, lower predicted OGT and pH, and a predicted competitor lifestyle would allow organisms to survive better in the soil ecosystem. Genomes from soil also encoded pathways beneficial to surviving in such ecosystem, e.g. N_2_ fixation, purine degradation to urea, and beta lactam resistance.

In conclusion, our results identify clear soil preferences for four classes (Terriglobia, Vicinamibacteria, Blastocatellia, and Thermoanaerobaculia) of Acidobacteriota. We also demonstrate that such preferences are driven not only by taxa previously recognized as prominent soil-dwellers in prior isolation and amplicon efforts, but also by multiple yet-uncultured orders and families in these four classes that are ubiquitous and abundant yet poorly characterized. Further, our analysis indicates that despite the observed preference patterns, most Acidobacteriota classes, orders, families, and genera are habitat generalists rather than specialists. As well, our global comparative genomic analysis provides new insights into the genomic features, predicted physiological optima, and metabolic repertoires of members of the Acidobacteriota, and disentangles the role played by phylogeny versus habitat in shaping Acidobacteriota genomic and predicted metabolic features.

## Acknowledgments

This work has been supported by the US National Science Foundation (NSF) grant 2016423 to N.H.Y. and M.S.E., and the US National Institute of Health (NIH) grant 1P20GM152333 to MSE.

